# Symmetric Fusion of fMRI and EEG for Spectrally Resolved Functional Neuroimaging

**DOI:** 10.64898/2026.01.31.703060

**Authors:** Jung-Hoon Kim, Zhongming Liu

**Affiliations:** Developing Brain Institute, Children’s National Hospital, Washington DC, USA; School of Medicine, George Washington University, Washington DC, USA; Department of Biomedical Engineering, University of Michigan, Ann Arbor, Michigan, USA; Department of Electrical Engineering and Computer Science, University of Michigan, Ann Arbor, Michigan, USA

**Keywords:** EEG, fMRI, multimodal recording, source imaging

## Abstract

Simultaneous electroencephalography (EEG) and functional MRI (fMRI) offers complementary sensitivity to fast electrophysiological dynamics of EEG and spatially resolved hemodynamics of fMRI, yet previous joint-analysis approaches are confined to fixed task paradigms and struggle with continuous or naturalistic brain states. We FSINC (Fusing Source Imaging based on a Neurovascular Coupling) model, a unified EEG–fMRI source imaging framework that reconstructs cortical activity to simultaneously explain both modalities. FSINC integrates frequency-resolved EEG source activity with fMRI via a data-driven neurovascular coupling model that estimates band-specific coupling coefficients (β) and accommodates a tunable spatial–temporal trade-off through hyperparameters (*λ*_2_, *λ*_3_). In realistic simulations, FSINC outperformed conventional methods (wMNE, LORETA) in both spatial and temporal accuracy across EEG SNRs (−10 to 10dB) and numbers of concurrent sources (up to five), with optimal performance at *λ*_2_ = 10^2^ and *λ*_3_=1 (e.g., LE: 0.51±0.24mm; SDI: 0.03±0.37mm; temporal accuracy: 0.95 ± 0.05). Applied to simultaneous EEG–fMRI during contrast-reversing visual stimulation (=5.95Hz), FSINC revealed stimulus-locked responses localized to early visual cortex and stimulus-induced modulation of intrinsic alpha oscillations extending into visual and attention networks, patterns that conventional methods failed to capture. Estimated β-weights were broadly consistent with prior reports of negative (theta/alpha) and positive (gamma) BOLD–electrophysiology associations. These findings demonstrate that FSINC enables high–spatiotemporal-resolution source imaging from EEG–fMRI recordings via data-driven hemodynamic modelling, and is expected to be well-suited for continuous and naturalistic brain states (e.g., resting state, natural moving-watching, and narrative listening) that are difficult to interrogate with either modality alone.

## 1. INTRODUCTION

Functional magnetic resonance imaging (fMRI) and electroencephalography (EEG) are two widely used neuroimaging modalities for non-invasive studies of human brain function. EEG measures neurogenic electrical potentials on the head surface with excellent temporal resolution, but poor spatial resolution (de Peralta-Menendez and Gonzalez-Andino, 1998; Michel et al., 2004; He et al., 2018). FMRI measures hemodynamic responses accompanying neural activity with relatively high spatial resolution, but limited temporal resolution (Bandettini et al., 1992; Kwong et al., 1992; Ogawa et al., 1992). Given their complementary merits and limitations, efforts have been made to integrate fMRI and EEG, or magnetoencephalography (MEG), for better resolution and specificity in both space and time (Liu et al., 1998; Dale et al., 2000; Portas et al., 2000; Laufs et al., 2003a; Laufs et al., 2003b; Martınez-Montes et al., 2004; Babiloni et al., 2005; Eichele et al., 2005; Debener et al., 2006; Goncalves et al., 2006; Im et al., 2006; Bénar et al., 2007; Mantini et al., 2007; Liu and He, 2008; Calhoun et al., 2009; Liu et al., 2013).

Existing approaches for fMRI-EEG integration fall into two categories (Jorge et al., 2014). In one category, namely fMRI-constrained EEG source imaging, the activation patterns obtained from fMRI data are used to constrain and improve EEG source imaging (Liu et al., 1998; Henson et al., 2010; Quirós et al., 2010; Murta et al., 2015). In the other category, namely EEG-informed fMRI mapping, features of EEG data are used to guide mapping of fMRI data (Thornton et al., 2011; Fahoum et al., 2012). Both strategies are asymmetric in the sense that results are first obtained from one modality and then used in the analysis for the other modality. Applications of these strategies have either improved the spatial resolution of imaging electrophysiological activity or addressed the neural correlate of fMRI activity. However, such asymmetric strategies fail to exploit the fundamental relationships between electrophysiology and hemodynamics.

In fact, the underlying sources of EEG and fMRI are tightly coupled. Both fMRI and EEG signals primarily result from synaptic inputs to neurons (Logothetis et al., 2001; Logothetis, 2002). Synaptic activity generates electrical potentials on the scalp surface (Mathiesen et al., 1998; Arthurs and Boniface, 2002), while the head acts as a spatially low-pass filter and severely reduces the spatial resolution and specificity of EEG (Michel et al., 2004). Synaptic activity also causes local metabolic and hemodynamic changes observable with fMRI sensitized to blood oxygenation level dependent (BOLD) contrast (Metea and Newman, 2006; Buxton, 2009). Neurovascular coupling acts as a temporally low-pass filter, thus limiting the temporal resolution and specificity of fMRI. Therefore, fMRI and EEG signals share, at least in part, a common origin that manifests itself in highly distinct spatial and temporal scales (He et al., 2018).

This notion is supported by the evidence obtained with simultaneously recorded neural and fMRI signals (Logothetis et al., 2001; Arthurs and Boniface, 2002; Goense and Logothetis, 2008). The BOLD fMRI signal accompanies, but lags behind, changes in local field potentials (LFP) that reflect synaptic input to neuronal ensembles (Buzsáki et al., 2012). The LFP-fMRI coupling is not confined to a single frequency band, but applies to many, if not all, frequency components of neural activity (Goense and Logothetis, 2008). The fMRI signal is also coupled with electrocorticography (ECoG) and EEG in a similar fashion (Goldman et al., 2002; Mukamel et al., 2005; Niessing et al., 2005; Goncalves et al., 2006; Wan et al., 2006; Yuan et al., 2010). However, different frequency bands of LFP, EEG, or ECoG contribute differently to the fMRI signal (Mantini et al., 2007; Conner et al., 2011; Liu et al., 2013). Disentangling different neuroelectric sources that contribute to the fMRI signal is also an “inverse problem”, which shares the ill-posed nature as the inverse problem of EEG (or MEG). The fMRI inverse problem applies to the time domain, whereas the EEG inverse problem applies to the spatial domain.

To integrate fMRI and EEG, we present herein a method to solve the inverse problems of fMRI and EEG altogether, unlike most previously reported asymmetric strategies. Central to this method is our assumption that the fMRI signal is linearly coupled to the power fluctuations of the frequency components of synaptic current that also generate EEG. Given this assumption, current sources can be estimated through an iterative algorithm in order to fit both fMRI and EEG data (Fig. 1). To evaluate its efficacy, we have tested this algorithm both in computer simulation and with data collected from humans under visual stimulation.

**Fig 1.**
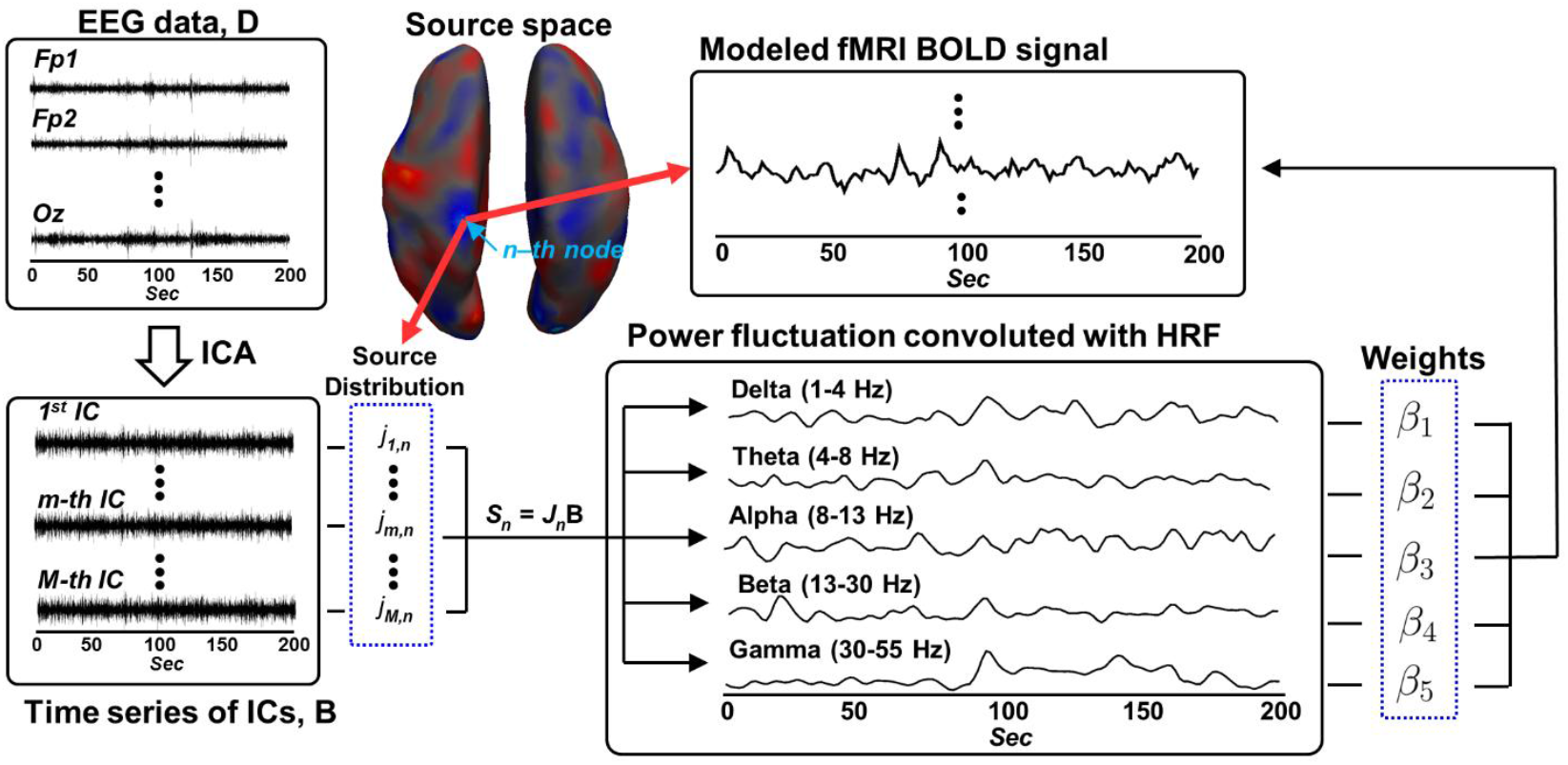
Schematic diagram of the proposed neurovascular coupling model. The black box represents variables driven from data whereas the blue dot box represents unknown variables in the model.

## 2. METHOD AND MATERIALS

For the notations in this paper, non-bold, italic, and lowercase letters are scalars; bold, italic, and lowercase letters are vectors; bold, regular and capital letters are matrices.

### 2.1. EEG inverse problem

For EEG, the forward problem concerns the mapping from a distribution of cortical current density to a distribution of scalp potentials. Reversely, the inverse problem concerns the mapping from scalp potentials to cortical current density. The forward problem is well determined and can be solved based on models of the cortex and the head with realistic geometries.

In this study, cortical current density was modeled with about 25,000 current dipoles evenly placed with ∼3mm spacing on the cortical surface generated from T_1_-weighted MRI (Fischl, 2012). Every dipole was assigned a fixed orientation perpendicular to the local cortical patch. To model the spatial mapping from current dipoles to scalp potentials, a boundary element method was used to calculate the lead-field matrix based on a piecewise homogeneous head model (Hamalainen and Sarvas, 1989). The model consisted of three compartments, i.e. the scalp, skull, and brain, with electrical conductivity of 0.33, 0.0041, and 0.33 S/m, respectively.

Through the lead-field matrix (**L** ∈ ℝ^*M*×*N*^), EEG data (**X** ∈ ℝ^*M*×*T*^) is expressed as a linear function of current sources (**S** ∈ ℝ^*N*×*T*^) plus noise (**σ** ∈ ℝ^*M*×*T*^), as defined in Eq. (1).

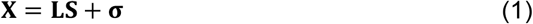

where *M, N*, and *T* are the numbers of channels, sources, and time points of EEG, respectively.

With EEG alone, the inverse problem is to estimate **S** given **X** and **L**. Since *M* ≪ *N*, this problem is ill-posed such that the inverse solver for mapping **X** to **S** cannot be uniquely determined. Regularization is required to impose constraints to the desired solution. The regularization depends on data and noise characteristics at each time point; accordingly, the optimal inverse solver is also time varying.

Instead of optimizing the inverse solver separately for each time point, we factorize the time-varying inverse solver in terms of a set of time-invariant inverse solvers. Firstly, independent component analysis (ICA) is used to decompose EEG into temporally independent components (ICs). Each component is expressed as the product of a spatial map (a time-invariant factor) and a time series (a space-invariant factor). In a matrix notation, **X** can be expressed as the product of spatial maps **C** and their corresponding time series **B** as in Eq. (2).

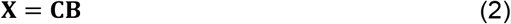

where **C** ∈ ℝ^*M*×*M*^, **B** ∈ ℝ^*M*×*T*^. Each column in **C** is a scalp map, and each row in **B** is a time series. Different rows in **B** are temporally independent and orthogonal, implying that **BB**^T^ = **I**. In Eq. (2), **C** and **B** can be solved given **X**, e.g. by the Infomax ICA algorithm (Lee et al., 1999).

The solution to Eq. (2) can be used to factorize the unknown source activity, since the linear mapping, **L**, from cortical sources to scalp potentials, is time invariant. Thus, **S** can be expressed as the product of the unknown source maps **J** and the known time series **B** as in Eq. (3), where **J** ∈ ℝ^*N*×*M*^.

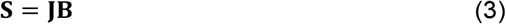

where each column in **J** is an unknown source distribution with the temporal dynamics expressed as the corresponding row in **B**.

By plugging Eqs. (2) and (3) in Eq. (1), and further multiplying **B**^T^ (i.e. the transpose of **B**) on both sides, Eq. (1) can be rewritten as Eq. (4).

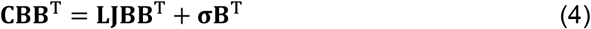

Since **BB**^T^ = **I**, Eq. (4) is simplified to Eq. (5)

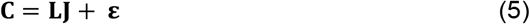

where the noise term **ε** = **σB**^T^.

In Eq. (5), each column in **J** is the source pattern underlying each corresponding column in **C**. Although it appears in a similar form as Eq. (1), Eq. (5) describes the EEG forward problem in terms of the spatial distributions of sources and data for every IC, as opposed to every time points.

It is worth noting that solving Eq. (5) is much more tractable than solving Eq. (1) for several reasons. First, the inverse solver only needs to be optimized for each component, instead of each time. As the number of ICs is orders of magnitude smaller than the number of time points, Eq. (5) requires much less time to solve. Secondly, the spatial pattern of each IC, as represented by the columns in **C**, appears more physiologically plausible and often easier to interpret than the spatial pattern at each time point. The instantaneous distribution of scalp potentials reflects a mixture of entangled source or noise patterns. Applying ICA separates the contributions from different source patterns of temporally independent dynamics. Moreover, the spatial patterns obtained by ICA often appear “dipolar” or focal yet distinctive across components. Therefore, it is more likely to arrive at an optimal solution for each component than for each time. Thirdly, ICA also helps to isolate and remove some noisy components (McKeown et al., 1998).

Importantly, after solving the inverse problem separately for every component, the inverse solution for every time point is readily derivable given Eq. (3). The time-variant source activity is thus factorized and expressed as the product of time-invariant spatial patterns (in the source space) and their corresponding temporal dynamics (derived from data in the sensor space).

### 2.2. fMRI inverse problem

Using the strategy elaborated in the previous subsection, one may already attempt to solve the EEG inverse problem based on Eq. (5) and Eq. (3). Since our focus is on the integrated analysis of fMRI and EEG, we choose to pair the EEG inverse problem with the fMRI inverse problem and devise a joint inverse solver to tackle both problems altogether.

To formulate the fMRI inverse problem, let us express the fMRI signal in terms of current source activity. Let *y*_*n*_(*t*) be the fMRI signal at the *n*-th cortical location and *S*_*n*_(*t*) be the source activity at the same location. Here, *S*_*n*_(*t*) refers to the *n*-th row and the *t*-th column of **S** as in Eq. (1) and Eq. (3). As a key assumption in our approach, the fMRI signal is modeled as the weighted sum of the power fluctuation of every frequency component of *S*_*n*_(*t*) after the power is temporally convolved with the hemodynamic response function (HRF) – a canonical model of neurovascular coupling (Logothetis et al., 2001).

Specifically, a frequency component of a signal, whether it is an unknown time-series of a source or a known time-series signal measured from a sensor, is defined or derived as a result of applying a bandpass filter to the signal. For simplicity, we refer to different frequency bands only by their center frequencies, {*f*_*p*_|*p* = 1 … *P*} without explicitly specifying their bandwidth. Of the source activity, *S*_*n*_(*t*), the *p*-th frequency component, *S*_*n*_(*p, t*), is defined as Eq. (6).

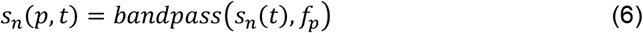

Given *S*_*n*_(*p, t*), 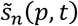 can be derived by the Hilbert transform.

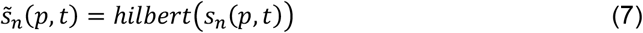

Given both *S*_*n*_(*p, t*) and 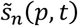, the band-limited power of *S*_*n*_(*p, t*) is expressed as Eq. (8).

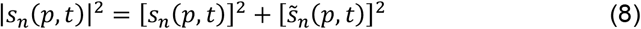

The fMRI signal is modeled as the weighted sum of the HRF-convolved power envelope across all frequency bands, as Eq. (9).

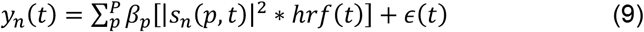

where *ϵ*(*t*) is the noise unexplained by the model. The weights ***β*** = [*β*_1_ … *β*_*p*_ … *β*_*P*_] indicate the relative contributions from individual frequency components to the fMRI signal.

Given Eq. (9), the fMRI inverse problem is formulated as the estimation of |*S*_*n*_(*p, t*)| for *p* = 1 … *P* given *y*_*n*_(*t*) for each cortical location *n*. This problem has no unique solution without considering any additional constraint, which is a dilemma similar to the EEG inverse problem.

### 2.3. Joint Inverse Problems of EEG and fMRI

Here, we approach both inverse problems together by utilizing the complementary spatial and temporal information from fMRI and EEG. We assume that neuroelectric activity is spatially transformed to give rise to EEG through linear volume conduction, and that its power fluctuations of band-limited components are HRF-convolved and summed to give rise to the local fMRI signal through neurovascular coupling. By solving both inverse problems altogether, the source estimate should fit both fMRI and EEG. The temporal information in EEG helps solve the fMRI inverse problem, and the spatial information in fMRI helps solve the EEG inverse problem. Following this rationale, we formulate the joint EEG-fMRI inverse problem as below.

Given Eq. (3), *S*_*n*_(*t*) can be written as a weighted sum of the independent time series in **B**.

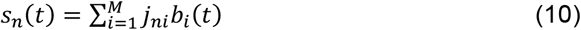

Since band-pass filtering is linear, *S*_*n*_(*p, t*) can be expressed as Eq. (11).

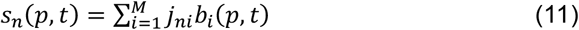

From Eq. (8), the band-limited power of *S*_*n*_(*p, t*) is further expressed as Eq. (12).

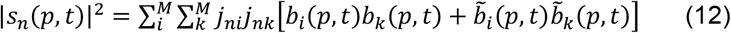

For the simplicity in notation, let us denote

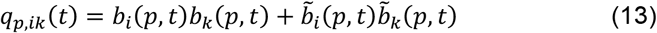

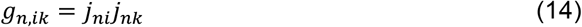

Let us further define two vector notations, ***g***_*n*_ and ***q***_*p*_(*t*), as below.

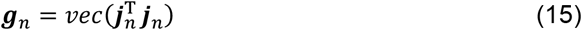

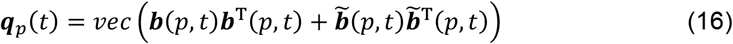

where 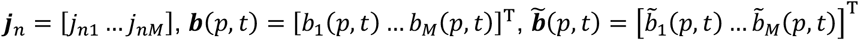, and *vec*(·) is a function that vectorizes a matrix into a column vector.

Then, Eq. (12) can be rewritten as Eq. (17)

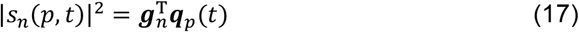

Note that Eq. (17) factorizes the band-limited power of source activity into a time-invariant term ***g***_*n*_ and a space-invariant term ***q***_*p*_(*t*). Note that ***q***_*p*_(*t*) is computable from EEG data by using ICA to obtain the IC time series **B** and then applying the bandpass filtering and Hilbert transform to the IC time series. As a result, ***q***_*p*_(*t*) provides a set of basis functions (i.e., time series). Their weighted sum is used to model the band-limited power of the source activity in the *p*-th frequency band. While the same basis set is applicable to every source location, the weights for the linear combination are variable across source locations. Note that ***g***_*n*_ depends only on the covariance of the sources underlying individual ICs of EEG, but independent of time and frequency. Given Eq. (17), Eq. (9) can be rewritten as Eq. (18)

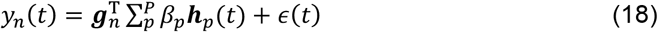

where ***h***_*p*_(*t*) = ***q***_*p*_(*t*) ∗ *hrf*(*t*), *hrf*(*t*) stands for the HRF, and ∗ stands for convolution.

Eq. (18) indicates that the fMRI signal at any given location is modeled as a weighted sum of known basis functions. The weights are factorized into a spatial factor ***g***_*n*_ and a spectral factor *β*_*p*_, which reflects the frequency-wise contribution to the fMRI signal at every source location. We can further rewrite Eq. (18) in two ways.

Given 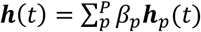, we rewrite Eq. (18) as Eq. (19)

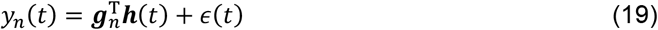

Given 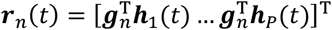, we rewrite Eq. (18) as Eq. (20)

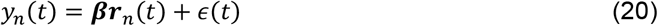

The joint inverse problem is to solve the unknowns in Eq. (5) and Eq. (18), such that the solutions fit both fMRI and EEG data. The error of fitting is defined in terms of the mean of squared errors, as Eq. (21) for EEG and as Eq. (22) or (23) for fMRI.

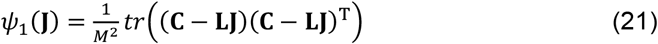

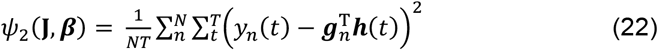

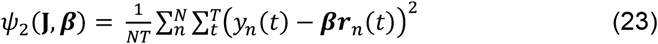

In addition, we add a third cost function. It is defined as the Laplacian-weighted norm of **J**. This term accounts for the physiological constraint that neighboring source activities tend to be correlated – an assumption also made in the Low-Resolution Brain Electromagnetic Tomography (LORETA) (Pascual-Marqui et al., 2002).

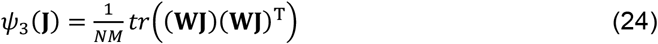

where **W** ∈ ℝ^*N*×*N*^ approximates the surface Laplacian operator (Wagner et al., 1996). Its elements {*w*_*ik*_} are defined as below.

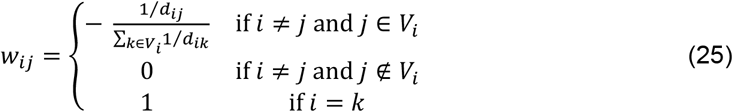

where *d*_*ij*_ is the cortical distance between the *i*-th source and the *j*-th source, and *V*_*i*_ is the set of indices corresponding to the neighbors of the *i*-th source, respectively.

The cost function is defined as the weighted sum of *ψ*_1_, *ψ*_2_, and *ψ*_3_, as in Eq. (26).

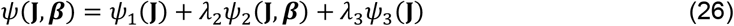

where *λ*_2_ and *λ*_3_ are hyperparameters that tune the balance the spatial- and temporal accuracies, respectively.

Note that this cost function only depends on two sets of unknowns: **J** and ***β***, because **g**_*n*_ is fully computable from **J** as shown in Eqs. (14) or (15).

To solve the joint inverse problems, we seek the optimal solution that minimizes the cost function.

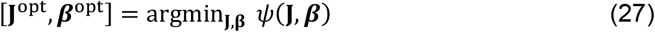

### 2.4. Solver of Algorithm

In the following, we describe a gradient descent algorithm to approach this optimization problem. For each term in the cost function, we derive its gradient with respect to **J** and ***β***. Since *ψ*_1_ and *ψ*_3_ are quadratic functions of **J**, their gradients with respect to **J** are simply linear functions, as Eq. (28) and Eq. (29).

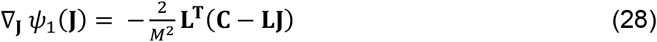

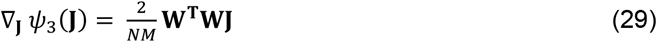

The second term in the cost function, *ψ*_2_, depends on both **J** and ***β***. Its partial derivative with respect to **J** and ***β*** is calculated iteratively such that the gradient at each step of iteration is calculated based on the iteratively updated estimates of **J** and ***β***, starting from their initial estimates.

The partial derivative with respect to each row of **J** (denoted as ***j***_*n*_) is expressed as Eq. (30).

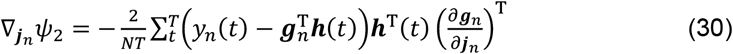

where 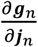 is the Jacobian matrix of ***g***_*n*_ with respect to ***j***_*n*_. It is *M*-by-*M*^2^, and its element indicates the partial derivative of each element in ***g***_*n*_ with respect to each element in ***j***_*n*_.

Let us define ***y***_*n*_ = [*y*_*n*_(1) … *y*_*n*_(*t*) … *y*_*n*_(*T*)] and **H** = [***h***(1) … ***h***(*t*) … ***h***(*T*)]. Eq. (30) can be rewritten with vector/matrix notations, as Eq. (31).

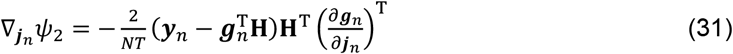

As shown in Eq. (31), this gradient is a non-linear function of ***j***_*n*_, implying that we cannot have an analytic form of solution that minimizes *ψ*_2_ without iteration.

It follows that the gradient of *ψ*_2_ with respect to **J** is written as Eq. (32).

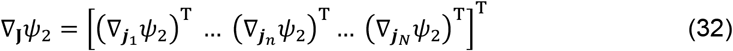

Combining Eqs. (28), (29), (32) yields Eq. (33)

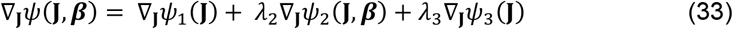

For the gradient of *ψ*_2_ with respect to ***β***, we start from Eq. (23) and define the followings.

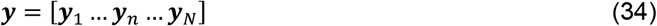

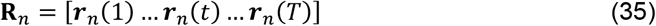

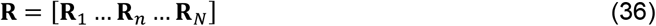

Then we can rewrite Eq. (23) as below

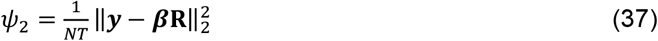

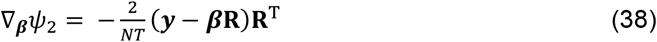

Note that ∇_**J**_*ψ*(**J, *β***) = ∇_***β***_*ψ*_2_, because ***β*** is only involved in *ψ*_2_. Eq. (38) is a linear function of ***β***, and therefore there is an analytic solution to ***β*** that minimizes *ψ*_2_ given any fixed **J** (and thus **R**). This solution is expressed as Eq. (39).

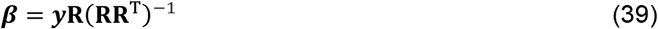

We implemented this algorithm in Matlab (MathWorks, Natick, USA), as described below (Fig 2).

**Fig 2.**
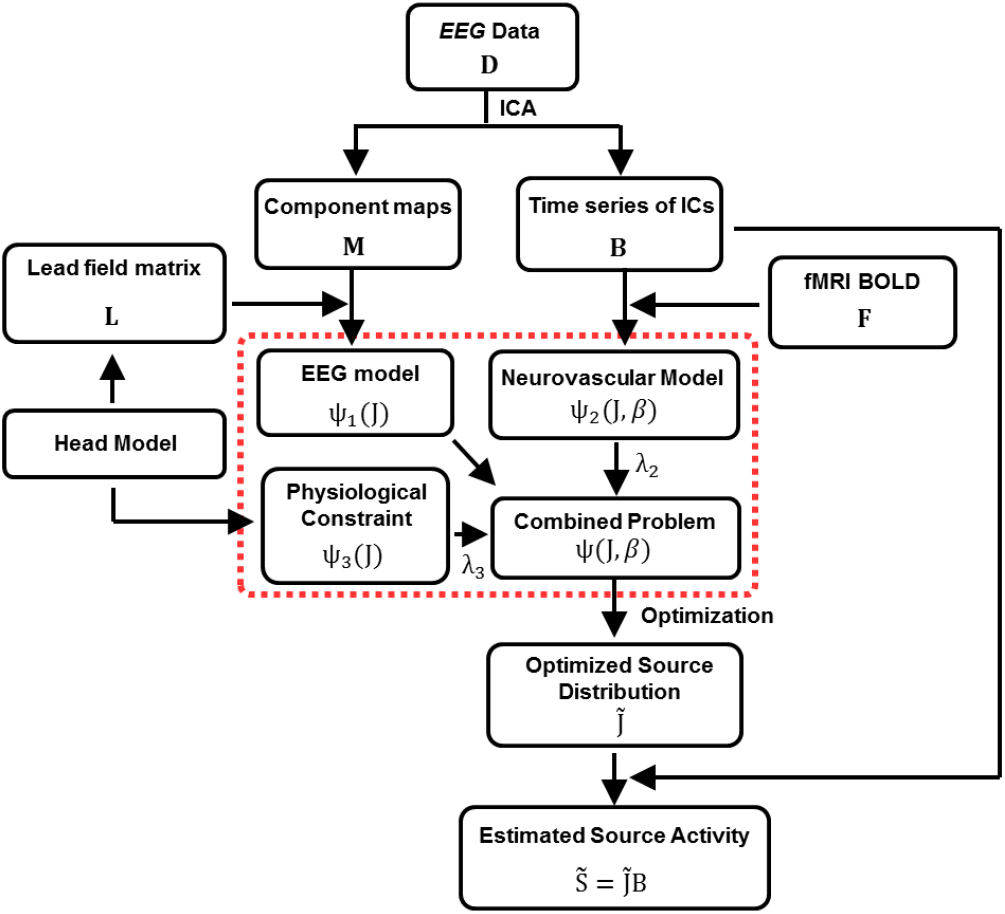
Diagram of the proposed source imaging method. Red box represents the unknown parameters that are estimated from the algorithm.

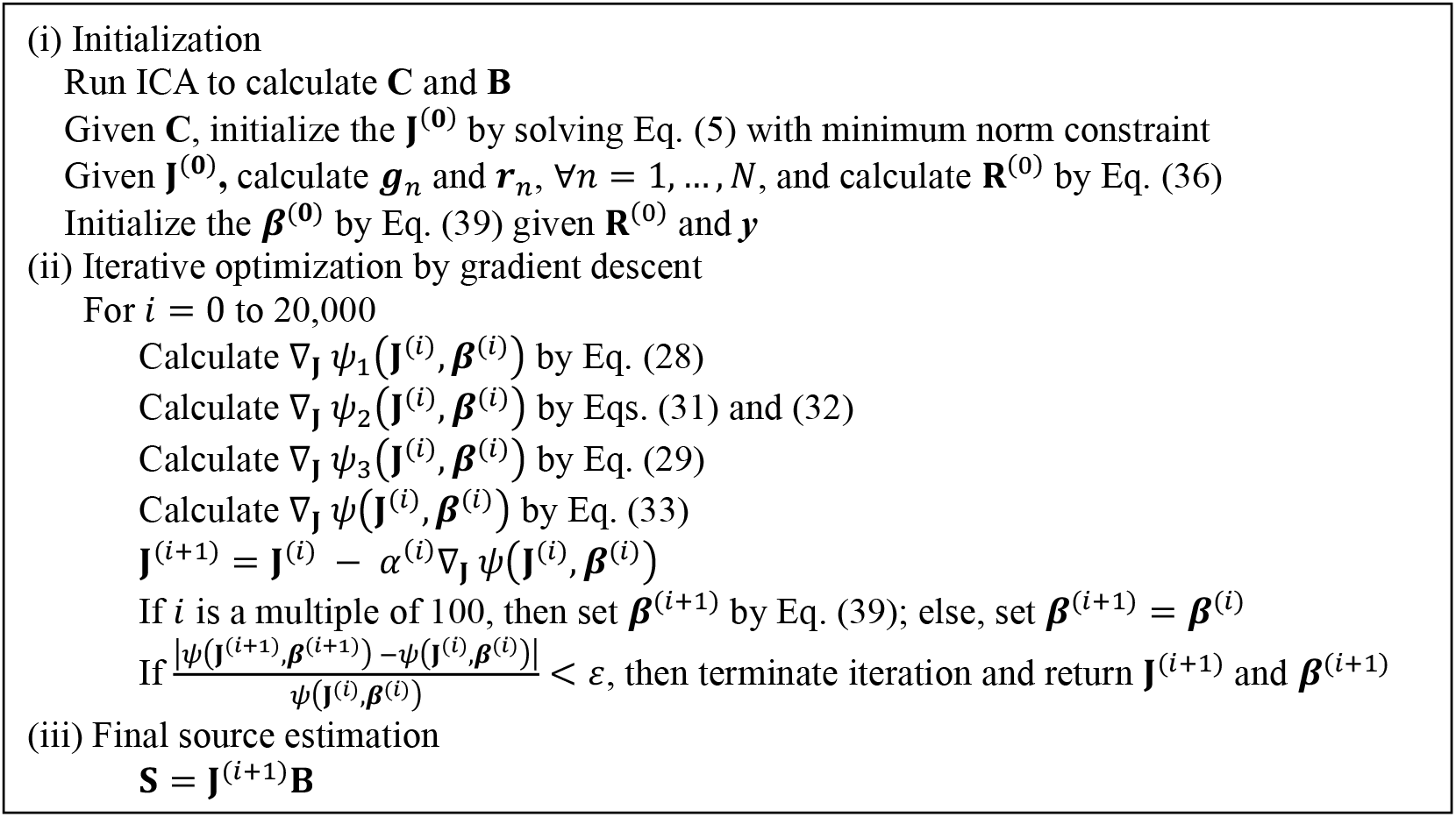

In this gradient descent algorithm, the step size, *α*^(*i*)^, was set by using the Barzilai-Borwein method (Barzilai and Borwein, 1988; Friedlander et al., 1998), as expressed in Eq. (40).

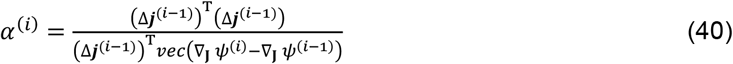

where Δ***j***^(*i*−1)^ = *vec*(**J**^(*i*)^ - **J**^(*i*−1)^). The stop criteria *ε* was set to 10^−10^.

### 2.5. Realistic simulations

We initially tested the proposed method with computer simulations. Multiple (from 2 to 5) extended sources were simulated with realistic cortical and head models. Each source extended from a randomly determined center to a 2D Gaussian cortical distribution with a standard deviation of 12 mm. Different sources were separated by a center-to-center distance of at least 30 mm. Every source was assigned with a distinct time series obtained by ICA of resting-state EEG data (acquired from a single subject) such that the simulated source activity was physiologically realistic. In this simulation, the source space included 24,673 cortical locations spaced by 3.19 mm; the simulated activity lasted about 3.5 min.

From the simulated sources, we simulated 32-channel EEG signals by using a realistic head volume conductor model by using FieldTrip (Oostenveld et al., 2011). To synthesize fMRI signals, source activity at each cortical location was filtered into five frequency bands (delta: 1-4 Hz, theta: 4-8 Hz, alpha: 8-14 Hz, beta: 14-31 Hz, gamma: 31-55 Hz). For every frequency band, the band-limited power was temporally convolved with the HRF. The resulting time series was combined across frequency bands through a weighted sum, where the frequency-wise weight was set to be [delta: −0.2, theta: −0.6, alpha: −0.6, beta: −0.3, gamma: 0.6], based on the finding from a previous study (Mukamel et al., 2005). White noise was simulated and added to the synthesized EEG and fMRI signals, giving rise to noise-contaminated EEG and fMRI data with the signal-to-noise ratio (SNR) equal to –10, –5, 0, 5, or 10 dB for EEG, and 5 dB for fMRI.

Specifically, simulation was performed in the following settings.

i. 2 sources, EEG SNR=5dB, fMRI SNR=5 dB, repeated for 5 times with source locations randomly determined for each time. The simulated data in this setting was used to test the effects of regularization parameters on imaging performance, and to further optimize the regularization parameters for the algorithm tested in the following two simulation settings as well as for the experimental data.
ii. 2 sources, EEG SNR = –10, –5, 0, 5, or 10 dB; fMRI SNR = 5 dB. Given each SNR setting, simulation was repeated for 40 times with source locations randomly determined for each time.
iii. 2, 3, 4, or 5 sources, EEG SNR=5 dB, fMRI SNR=5 dB. Simulation was repeated 20 times with source locations randomly determined for each time.

For each simulation, we estimated the source activity by using the proposed method. The spatiotemporal distribution of the estimated source activity was evaluated for accuracy in both space and time. To evaluate the spatial accuracy, we calculated an “activation” map, in which the activation value at each source location was the temporal standard deviation of the source activity estimated at that location. The estimated activation maps, the cortical locations with a value smaller than two times the standard deviation of source activity were fixed to zero. Given the thresholded activation map, we measured the localization error (LE) as the Euclidean distance between the center of the simulated source pattern μ and the center (of the mass) of the reconstructed source pattern 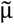. The center of reconstructed source patterns was defined as in Eq. (41)

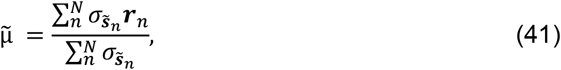

where r_n is the MNI coordinate of the n-th vertex. A higher LE indicated that the center of the estimated source activity was more distant from the predefined center, and vice versa. We further estimated the extent of the reconstructed source pattern, termed as source distribution index (SDI), as used in other papers, e.g., (Samadi et al., 2016).

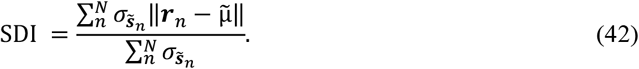

Note that the SDI of the actual activation map with a radius of 20 mm was about 8 mm, and the difference between SDI from actual source pattern and SDI from reconstructed source pattern was reported.

Temporal accuracy was calculated by averaging the correlation coefficients between the actual source activity and the estimated source activity at each cortical location. Considering computation capacity, temporal accuracy was estimated for a 25-seconds fragment of whole data (total 5000 samples). Here, a correlation of 1 meant the estimated source activity was identical to the actual source activity, a correlation of −1 indicated that the estimated source activity was identical to the actual source activity with an opposite phase, and a correlation of 0 indicated that there was no correlation between the actual source activity and the estimated source activity. For the simulated data set (i), the parameter setting (*λ*_2_ = 10^2^, *λ*_3_ = 1) was chosen to achieve the best spatiotemporal accuracy.

For comparison, we also did source imaging with alternative methods: weighted minimum estimation norm (wMNE) method (Tikhonov et al., 1977) and LORETA (Pascual-Marqui et al., 2002), were applied to the simulation data set (ii) and (iii). Weighted MNE and LORETA were implemented by MATLAB scripts built in-house and their optimal regularization parameters were chosen through Tikhonov regularization strategy, implemented by the function, *l_curve*, provided from the MATLAB toolbox (Hansen, 2007).

### 2.6. Performance validation with real EEG-fMRI data

Further, we tested the imaging performance of the proposed source imaging method with simultaneously acquired fMRI and EEG data from a total of 13 healthy subjects in accordance with a protocol approved by the Institutional Review Board at the National Institutes of Health. Each subject was presented with full-field black-and-white checkerboard visual stimuli reversing at 5.95 Hz with a 30s-ON-30s-OFF block design paradigm, repeated three times.

From each subject, EEG and fMRI data were simultaneously acquired in a 3-tesla human MRI system (General Electric Healthcare, Milwaukee, USA). EEG signals were recorded by using a 32-channel MRI-compatible recording EEG system (BrainProducts, Germany) with a sampling rate of 5 kHz and the analog bandwidth from 0.1 to 250 Hz. Blood oxygenation level dependent (BOLD) fMRI images were acquired by using a 16-channel receive-only phase array coils and a single-shot gradient-echo echo-planar imaging sequence (90° flip angle, 30 axial slices, 4 mm slice thickness, 30 ms echo time, 1.5 s repetition time, 220×165 mm^2^ field of view, matrix size =64×48, and sensitivity-encoded acceleration at a rate of 2). T1-weighted MRI was also acquired with a 3-D magnetization-prepared rapid gradient-echo sequence (12° flip angle, 725 ms inversion time, 2.25 ms echo time, 5 ms repetition time, and 1 mm^3^ isotropic voxel size).

From the recorded EEG data, gradient artifacts and cardiac ballistic artifacts were removed by using an algorithm as previously described (Liu et al., 2012). Then the data were resampled to 250 Hz and re-referenced to the average across channels. ICA was further applied by using EEGLAB (Delorme and Makeig, 2004). We used already preprocessed BOLD fMRI with standard preprocessing procedures, e.g., similar to ones described (Chang et al., 2013). In this study, we further detrended the preprocessed BOLD fMRI signal by regressing out third-order polynomials and then converted to percentage change separately for each voxel.

To obtain the fMRI activation maps in response to the visual stimulation, we computed the correlation between the vertex-wise fMRI time series and the box car of the visual paradigm convolved with the canonical HRF (cHRF). For EEG-derived source activity at the stimulus frequency (SF, 5.95 Hz), band-limited power fluctuations were computed and averaged in 1 s windows with 50% overlap. SF activation maps were generated by calculating the Pearson correlation between the cHRF-convolved boxcar and the SF band-limited power time course. For induced alpha activity (IAF, 8–14 Hz), we first removed harmonics of the SF from the source activity, then computed band-limited power, convolved it with the cHRF, and downsampled to the BOLD sampling rate. The IAF activation map was obtained via vertex-wise partial correlation between the BOLD signal and the convolved IAF power, controlling for the SF component. Subject-level correlation coefficients were projected to standard MNI space and transformed to z-scores using Fisher’s r-to-z transformation. Standardized maps were averaged across subjects, and statistical significance of the mean z-scores at each vertex/voxel was assessed using one-sample t-tests. Multiple-comparisons correction was applied to the BOLD and SF maps using false discovery rate (FDR) control at q < 0.05. IAF maps were evaluated without multiple-comparisons correction (p < 0.05). For visualization, the averaged z-scores were transformed back to r-values.

## 3. Results

We systematically compared the proposed FSINC algorithm with two conventional EEG source imaging methods, wMNE and LORETA, using simulated data. In the simulations, wMNE and LORETA could not be applied to the fMRI component because they require a prior statistical fMRI map; by contrast, FSINC directly integrates the fMRI time series and does not rely on such priors. In the experimental visual-task EEG-fMRI dataset, fMRI-derived information was incorporated for all three approaches, but differently, via direct time-series fusion for FSINC and via fMRI-informed priors for wMNE and LORETA.

### 3.1. Performance validation of proposed FSINC on simulation data

First, we searched the optimal parameter settings for the FSINC using the simulated dataset. Using the simulated dataset (i), we identified parameter values that jointly optimized spatial and temporal accuracy. A clear trade-off emerged across parameter settings: configurations with low *λ*_2_and a high *λ*_3_ performed like LORETA (Fig. 3, bottom left), whereas configurations with high *λ*_2_ and low *λ*_3_ produced the opposite pattern, higher spatial accuracy but lower temporal accuracy (Fig. 3, bottom right). Among the different parameter settings, the parameter setting with *λ*_2_ = 10^2^ and *λ*_3_ = 1 showed the best performance in both temporal and spatial aspects (LE: 0.51 ± 0.24 mm, SDI: 0.03 ± 0.37 mm, and temporal accuracy: 0.95 ± 0.05).

**Fig 3.**
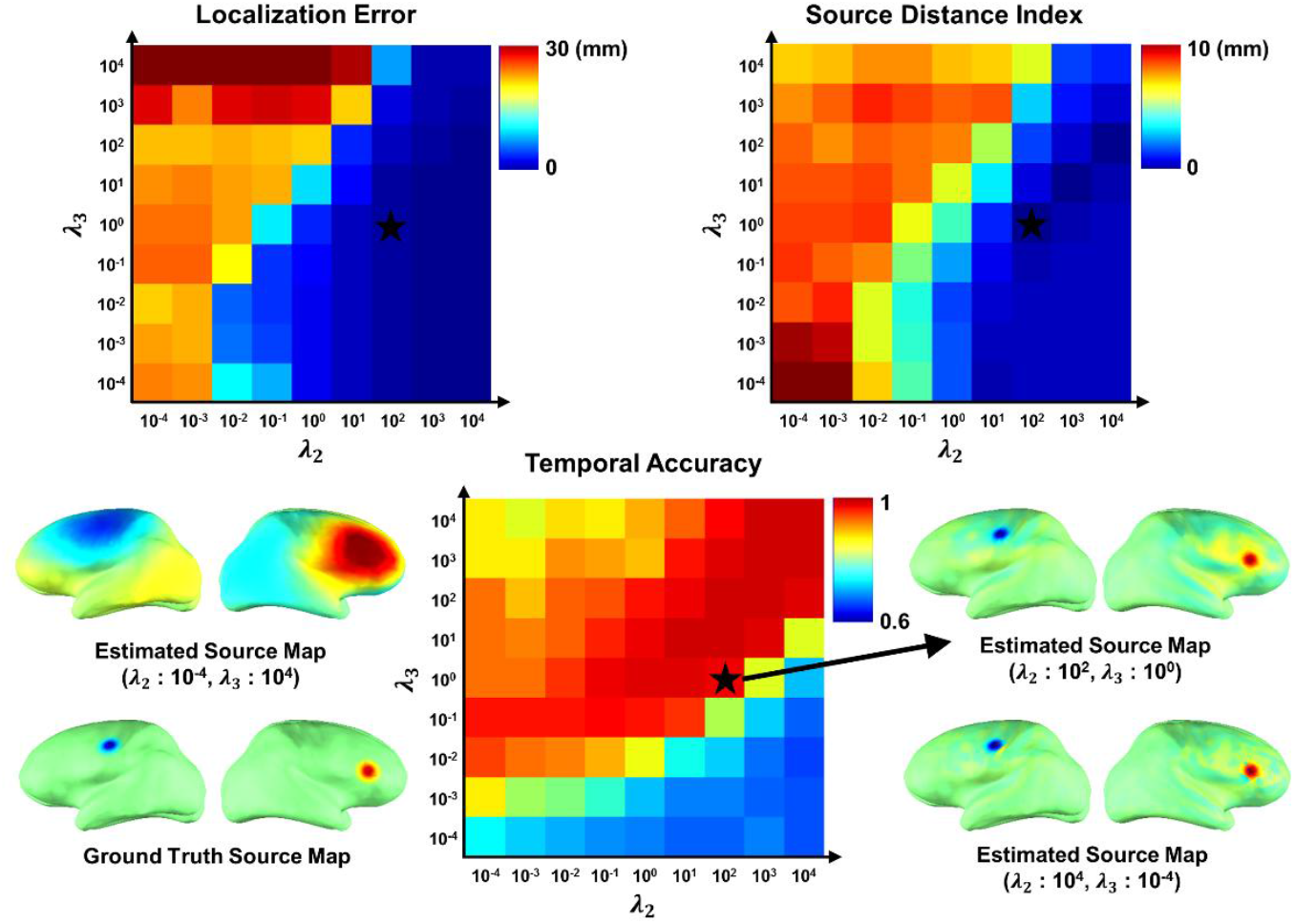
Performance comparison for the different parameters in the simulation data. The black star marks the parameter set that achieved the best overall performance in the simulation and was selected for subsequent analyses. EEG and fMRI SNRs were both set to 5 dB.

We then evaluated this selected parameter set across EEG SNR levels from −10 to 10 dB (with fMRI SNR fixed at 5 dB). Across all SNRs, wMNE outperformed LORETA on LE but underperformed on SDI (Fig. 4). Although the temporal accuracy of both methods improved with increasing SNR, neither recovered accurate spatial distributions across the full SNR range. By contrast, FSINC consistently outperformed both methods, spatially and temporally, at all SNR levels and remained robust even at the lowest SNR tested (−10 dB).

**Fig 4.**
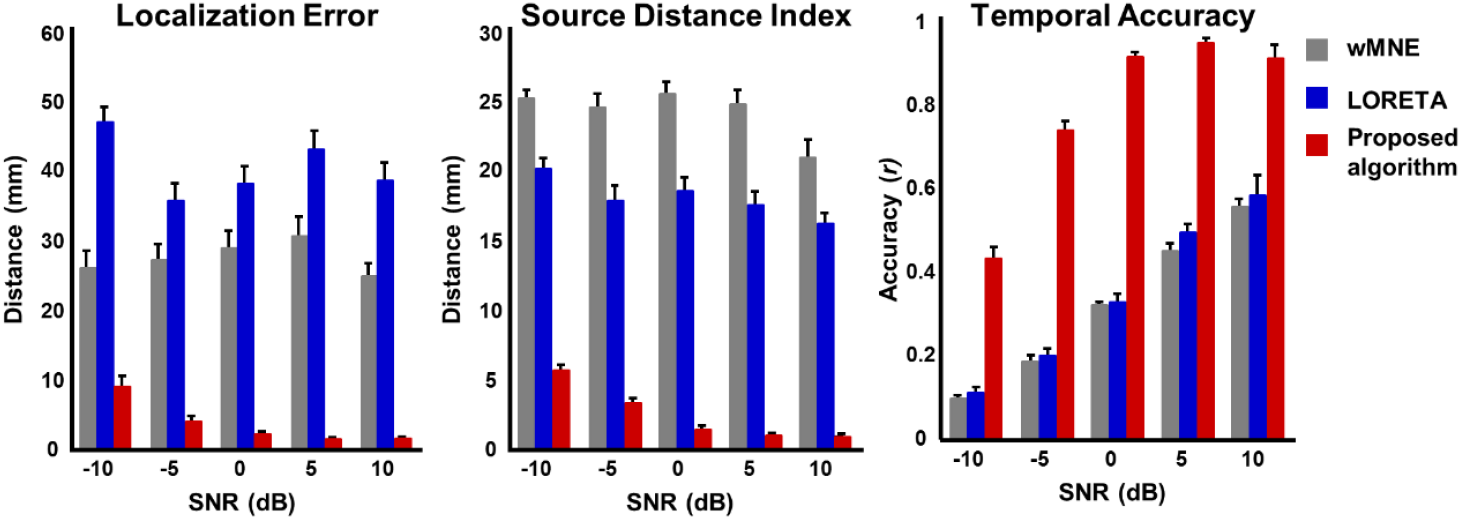
Performance comparison for the different SNR scale in the simulation data. Error bars denote standard error. The fMRI SNR was fixed at 5 dB. Each color stands for different source imaging methods.

We further assessed robustness with varying numbers of sources (up to five). As shown in Fig. 5, FSINC maintained similar spatial and temporal accuracy across source counts and performed favorably relative to wMNE and LORETA. Altogether, the simulation results indicate that FSINC outperforms conventional methods in both spatial and temporal accuracy.

**Fig 5.**
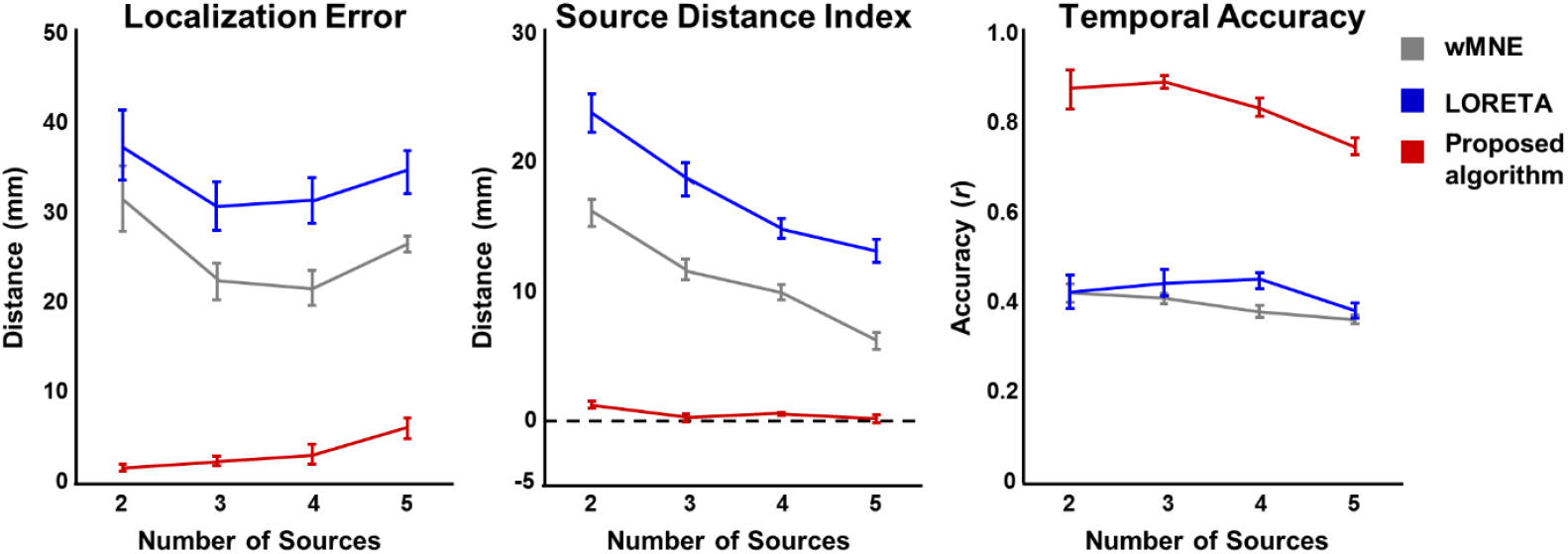
Performance comparison for the different number of source maps in the simulation data. The error bar indicates standard error. The SNR level of the fMRI is set to 5 dB. Each color stands for different source imaging methods. Error bars stand for standard error of mean.

### 3.2. FSINC method outperforms other source imaging methods in experimental data

Based on the superior performance on simulation data, we validated FSINC on the experimental dataset. We applied FSINC to the simultaneous EEG–fMRI recordings to estimate source images with high spatiotemporal resolution and to compute frequency-band weights. After normalizing per subject and averaging across subjects, the weights for the delta, theta, alpha, beta, and gamma bands were −0.08± 0.06, 0.37± 0.09, −0.21±0.15, −0.13±0.14, and 0.18±0.13, respectively (Fig. 6).

**Fig 6.**
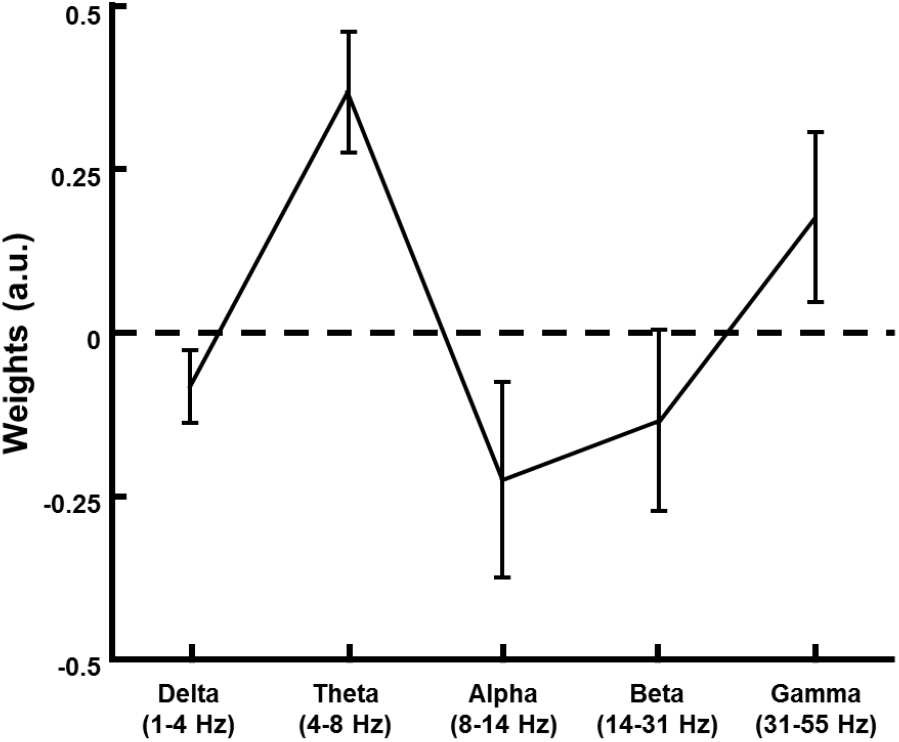
Mean optimized weights for each frequency band. Weights were normalized within subjects and averaged across subjects. Error bars denote standard error of mean.

Next, We next compared BOLD-activated regions with SF activation maps (sensor- and source-level) derived using different source imaging methods, at both single-subject and group levels. At the single-subject level (Fig. 7), the BOLD signal showed a strong positive correlation in the primary visual cortex, alongside widespread negative correlations extending beyond visual areas (Fig. 7A). The sensor-level SF map exhibited strong positive correlations over parietal and frontocentral channels, without negative correlations (Fig. 7B). At the source level, all methods showed positive activation in parietal regions that included visual cortex. However, wMNE failed to capture the early visual cortex in deeper (i.e., more inferior/posterior) locations. LORETA localized activity in deeper regions but did not encompass the early visual cortex and showed limited spatial specificity. In contrast, FSINC selectively covered the early visual cortex (Fig. 7C), closely resembling the positive BOLD activation pattern, while not showing activation in regions where the BOLD response exhibited negative correlations (Fig. 7A,C).

**Fig 7.**
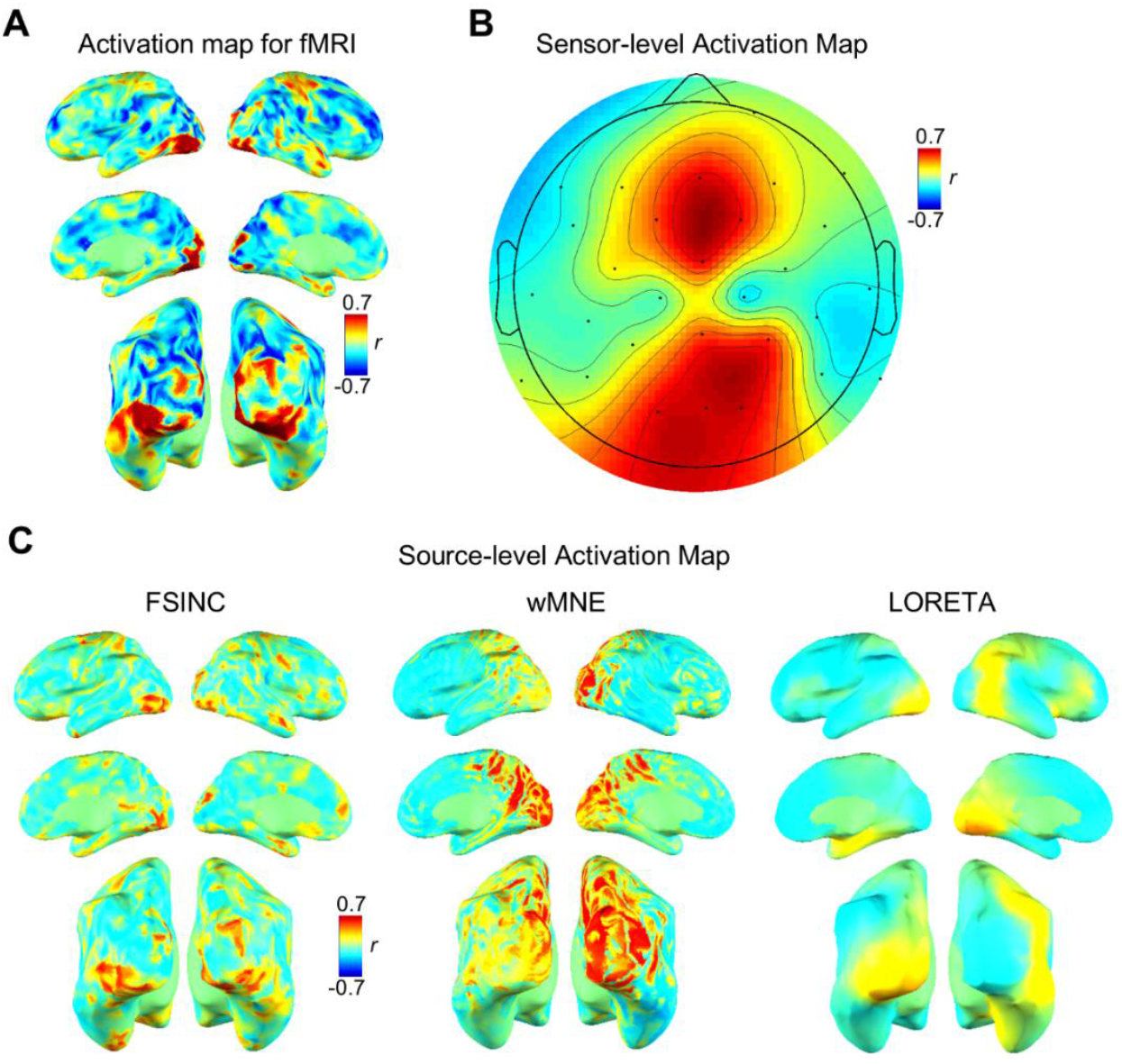
Single-subject activation comparisons. (**A**) correlation coefficient maps between the fMRI BOLD signal and the paradigm convoluted with the hemodynamic response function. (**B**) Correlation coefficient maps between the paradigm and the band-limited power at stimulation frequency (5.95 Hz) at the sensor level. (**C**) Same as (**B**), at the source level.

At the group level (Fig. 8), results mirrored the single-subject findings. wMNE yielded significant activation predominantly in more superficial visual regions (i.e., geometrically closer to sensors), while LORETA did not produce a statistically significant SF activation map. Interestingly, both the group-level BOLD map and the SF map derived with FSINC showed robust positive activation in the early visual cortex. For the IAF analysis, wMNE and LORETA showed significant correlations only in limited regions, whereas FSINC reconstructed an activation pattern extending into the extrastriate cortex, as well as frontal and parietal cortices, encompassing regions implicated in the attention network (Fig. 8, bottom). These findings suggest that the IAF and the BOLD response in the extrastriate cortex may arise from a shared spontaneous neuronal process not driven by the flickering visual task used in this study.

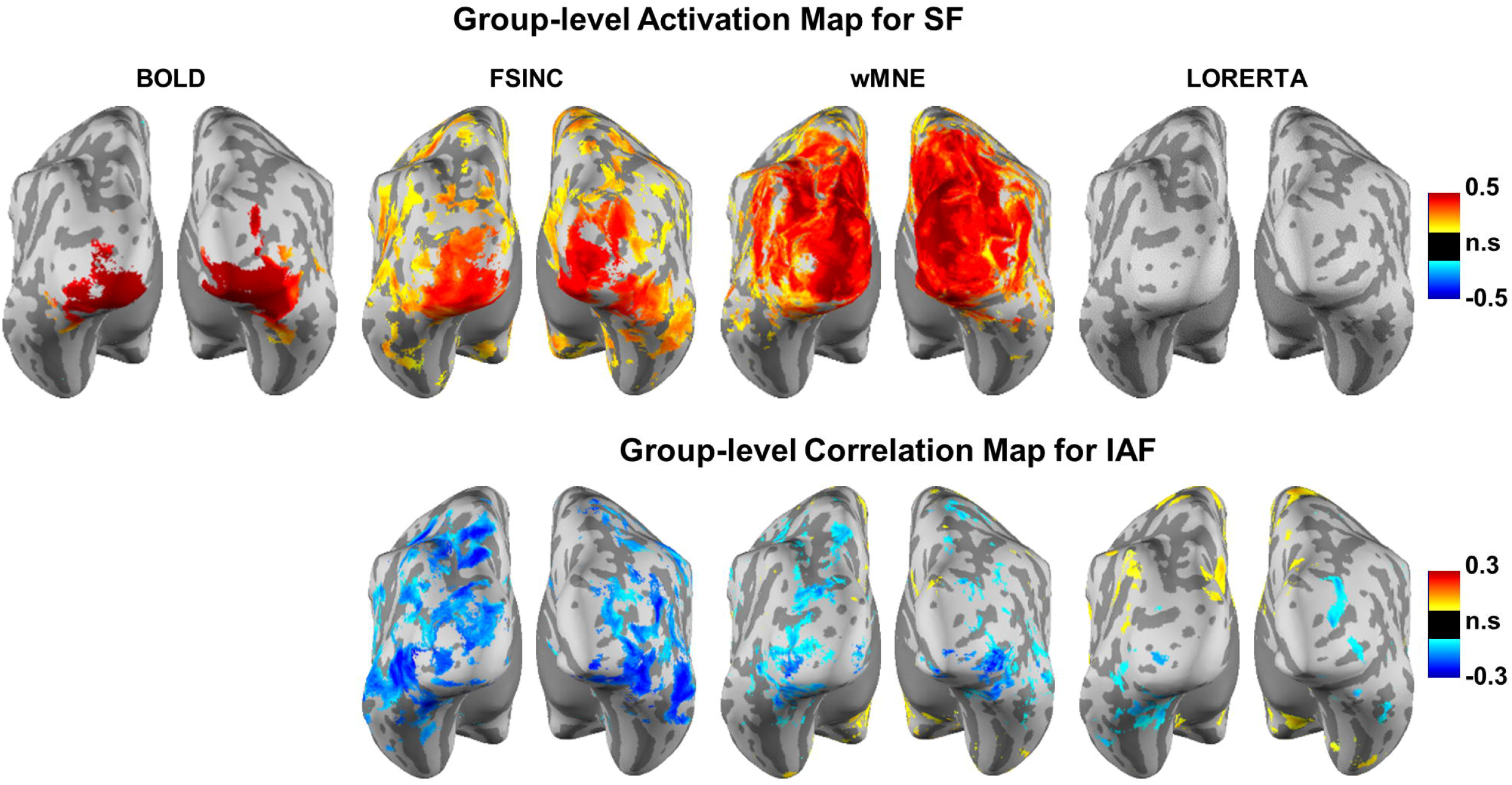

## 4. Discussion

In this study, we introduced FSINC, a data-driven EEG–fMRI integrated source imaging framework that models neurovascular coupling directly from band-limited power of source activity and fuses EEG and fMRI by incorporating an optimally estimated coupling model. Validation on both realistic simulations and experimental visual-stimulation data demonstrated that FSINC delivers superior spatial and temporal performance relative to conventional EEG source imaging methods (wMNE and LORETA) in simulations, and yields activation patterns that align closely with task-related BOLD responses while preserving EEG-level temporal resolution.

In the simulation study, we benchmarked FSINC against wMNE and LORETA. wMNE is a classical solution to the EEG inverse problem that compensates for depth bias via weighting, yet localization remains sensitive to anatomy and tends to under-recover deeper sources. LORETA augments the inverse solution with a spatial smoothness (Laplacian) constraint and depth weighting, favoring distributed, low-spatial-frequency solutions. These characteristics, well summarized in review work (Grech et al., 2008), explain their complementary strengths and weaknesses, as observed in our parameter searching experiment: settings with low λ_2_ and high λ_3_ reproduced LORETA-like behavior with higher temporal but lower spatial accuracy, whereas higher λ_2_ and lower λ_3_ favored spatial precision at a temporal cost. FSINC navigated this trade-off effectively; the setting λ_2_ = 10^2^, λ_3_ = 1 jointly optimized spatial and temporal accuracy in simulations. Critically, tuning λ_2_ and λ_3_ will provide a practical lever to balance spatial vs. temporal accuracy of estimated source activity for future applications.

Because fMRI’s temporal resolution is orders of magnitude lower than EEG’s, we leveraged independent component analysis (ICA) to obtain temporally independent EEG components with associated spatial maps, which can stabilize the mapping between electrophysiology and hemodynamics. Consistent with prior work (Makeig et al., 1996; Onton et al., 2006), a LORETA-like configuration (λ_2_ = 10^−4^, λ_3_ = 10^−1^) achieved improved temporal accuracy (0.70±0.10) compared with LORETA without ICA (0.49±0.13) at the same SNR, suggesting that incorporating ICA can improve temporal fidelity in general inverse pipelines.

Accumulating evidence indicates that BOLD responses relate differentially to frequency-specific electrophysiological activity: lower-frequency bands (e.g., theta/alpha) often show negative associations with BOLD, whereas higher-frequency (e.g., gamma) tends to show positive associations across regions, states, and species (Mukamel et al., 2005; Yuan et al., 2010; Murta et al., 2016). Such spectral-distinctive coupling was validated across different recording modalities (e.g. EEG, LFP, and EcoG) (Niessing et al., 2005; Khursheed et al., 2011; Scheeringa et al., 2011). Guided by this, FSINC estimates band-specific coupling (β) weights directly from data. We assumed these β weights to be spatially uniform, an approximation supported by reports of modest within-band regional variability (Conner et al., 2011). Our estimated weights (Fig. 6) were broadly consistent with prior findings, supporting the validity of the proposed FSINC approach. A notable deviation emerged in the theta band, which may reflect the task design: the flicker frequency (5.95 Hz) falls within the theta range (4–8 Hz), potentially altering cross-band interactions and the effective coupling in that band.

At the single-subject level (Fig. 7), SF-related activation localized to the primary visual cortex without mirroring regions that exhibited negative BOLD correlations to the stimulus. Negative BOLD responses during visual stimulation are well documented alongside positive occipital activations (Saad et al., 2001). Proposed mechanisms include neuronal inhibition and/or disproportionate metabolic demands (Shmuel et al., 2002), implying distinct neuronal origins from the positive BOLD network. Traditional fMRI-informed EEG inverses that rely on thresholded **p**-value maps treat positive and negative BOLD regions equivalently as “activated,” potentially conflating networks with different neurophysiological bases (Liu et al., 1998; Liu and He, 2008). In contrast, by learning the coupling from frequency-resolved EEG sources, FSINC can differentially weight frequency contributions and thus better separate networks with divergent neurovascular signatures—consistent with the observed alignment to positive BOLD in early visual cortex (Fig. 7A,C; Fig. 8). Moreover, our results recapitulate known functional dissociations: SF activity was confined to early visual cortex (Fig. 7C; Fig. 8), whereas IAF-related maps extended into dorsal/ventral attention systems, including extrastriate, frontal, and parietal cortices. This pattern echoes prior findings that alpha dynamics track attention and large-scale control networks (Di Russo et al., 2007; Vialatte et al., 2010), underscoring FSINC’s utility for probing ongoing neuronal processes during continuous, complex dynamics rather than only event-locked responses.

There are, however, a couple of limitations in this study. First, we selected hyperparameters based on simulation performance. In practice, optimal settings may vary with data quality (e.g., EEG/fMRI SNR) and user priorities along the spatial–temporal trade-off (Fig. 3). Future work should incorporate principled selection strategies, such as, nested cross-validation, empirical Bayes, or evidence optimization, and potentially adapt parameters per subject or even per region. Second, while a linear combination of band-limited powers explained a substantial fraction of BOLD variance, residual nonlinearities in neurovascular coupling likely remain (Siero et al., 2013). Extending FSINC to encompass nonlinear and state-dependent effects, potentially via kernelized regression, generalized additive models, or physiologically informed dynamic models (with subject-specific HRFs and regional delays), may further improve fidelity. Future works may include relaxing the spatial uniformity of β weights (with appropriate regularization), modeling cross-frequency interactions (e.g., phase–amplitude coupling), and testing generalization across tasks and resting-state paradigms.

## 5. Conclusion

Unlike prior approaches that either constrain EEG-based imaging with fMRI priors or use EEG to inform fMRI mapping, we introduced FSINC, a unified EEG–fMRI source imaging framework that reconstructs cortical activity to simultaneously account for both modalities. In realistic simulations, FSINC achieved superior spatial and temporal accuracy compared with conventional methods (wMNE, LORETA), by adjusting key hyperparameters (λ_2_ and λ_3_) implemented in FSINC. Applied to experimental visual-stimulation data, FSINC revealed stimulus-locked responses confined to early visual cortex and widespread, stimulus-induced modulations of intrinsic alpha oscillations across visual and attention networks; patterns that wMNE and LORETA failed to capture. Together, these findings demonstrate that joint EEG–fMRI inference can resolve fast, distributed neural dynamics with hemodynamic validation, opening new avenues for investigating continuous brain states that are difficult to access with either modality alone.

## Acknowledgment

This study was funded by the NIH MH104402. The funding sources had no involvement in the design, data collection, analysis, interpretation, writing of the report, and/or the decision to submit this article for publication. We would like to thank Drs. Yizhen Zhang and Kun-Han Lu for helpful discussion.

